# Regional species coexistence despite local priority effects: the overlooked role of dispersal–community feedback

**DOI:** 10.1101/2025.03.31.646410

**Authors:** Lucas A. Nell, Christopher A. Klausmeier, Tadashi Fukami

## Abstract

Many cases of animal-mediated dispersal are non-random, with the animals altering their movement pattern in response to the local species composition of the organisms that the vectoring animals disperse. Yet, this dispersal–community feedback has received little attention in metacommunity ecology. We use a mathematical model to show that dispersal–community feedback can promote regional species coexistence. As a well-characterized system, our model focuses on nectar-inhabiting bacteria and yeast that are dispersed by pollinators and affected by priority effects within flowers once dispersed. Model analysis suggests that bacteria and yeast coexist regionally across a wide range of regional pollinator abundances only when their occurrence in flowers influences the frequency of flower visits by pollinators. This microbe–pollinator feedback creates positive frequency dependence in each plant, causing competitive exclusion at the plant scale, but spatial partitioning across multiple plants, stabilizing coexistence at this scale. Our finding highlights dispersal–community feedback as an overlooked potential contributor to species coexistence.

**Significance Statement:** Many organisms rely on animals to move between habitats, and those animals often choose where to go based on which species are already there. We modeled yeast and bacteria that live in flower nectar and are carried between flowers by pollinators, which avoid nectar soured by bacteria. On a single plant, pollinators avoiding bacteria-laden nectar reinforces whichever microbe arrives first, so each plant becomes dominated by one microbe. Across many plants sharing a pool of pollinators, the same feedback lets both microbes persist over a wide range of pollinator abundances: yeast-dominated plants keep their pollinators and bacteria-dominated plants lose them, so each microbe has plants where it wins. Such dispersal–community feedback may be an overlooked mechanism of species coexistence.

## Introduction

Spatial scale plays a central role in promoting the coexistence of competing species in spatially structured environments (1–3). Conditions and interactions at local scales, including those causing exclusion and instability, can generate spatial heterogeneity that results in negative feedbacks and stable coexistence at larger scales (2, 4, 5). In environments where competitive rankings do not vary with space, local extinction–colonization events combined with appropriate life-history tradeoffs among competitors can cause regional coexistence (6–9). Alternatively, underlying spatial heterogeneity in the environment (e.g., resource types, predator abundance) can allow multiple competitors to specialize in the type of habitat that best suits them and therefore coexist (10–12).

Another way spatial heterogeneity can emerge is through local priority effects, where positive frequency-dependent competition causes alternative community states that depend on which competitor arrives first (13–18). However, it remains unclear whether coexistence at larger spatial scales can result from local alternative states. Because the probability of arrival to a new patch is likely to be proportional to a species’ regional abundance, this may generate positive feedback resulting in either competitive exclusion or alternative states at the regional scale instead of coexistence (19–21).

One possible solution to the problem of regional coexistence is non-random patterns of dispersal. Although most theoretical work on spatial coexistence assumes random dispersal, non-random dispersal has been shown to affect species coexistence in consumer-resource (22, 23) and competitive systems (24–27). Non-random dispersal is typically incorporated in models by including responses of competitors to environmental cues. Yet, many organisms (e.g., plants, microbes, pelagic larvae) have limited ability to alter their dispersal behavior based on the environment, especially over long distances or when suitable habitat patches are situated within an inhospitable matrix. In these dispersal-limited organisms, long-distance dispersal is often mediated by physical forces or by animals more mobile than themselves (28–30).

Animal-mediated dispersal is especially interesting in the context of coexistence (but see 31, for coexistence through a physical force) because of the non-random regional dispersal patterns they could generate for the species that they disperse (their “hitchhikers”). For example, clustered animal dispersal could cause aggregated patterns of hitchhiker abundances that could promote coexistence if it results in weakly or negatively correlated spatial distributions of competing hitchhikers (32–34). If animals preferentially go to habitats that are higher in quality for their hitchhikers, this could result in covariance between hitchhiker densities and growth rates, another mechanism predicted to promote coexistence (35).

Despite a relative lack of attention to animal-mediated dispersal affecting hitchhiker communities, empirical evidence suggests that it could be impactful in a variety of systems. Examples in ant-dispersed shrubs (36, 37) and sedges (38), as well as bird-dispersed mistletoes (39), show animal-mediated dispersal directing seeds to species-specific, high-quality habitats where they are less likely to be competitively excluded. For plants with herbivore-dispersed seeds, their dispersal trajectories are governed by the same organisms that consume them and likely respond to their presence and those of their competitors (29). Herbivores preferring areas with high abundances of the plants whose seeds they disperse could promote coexistence via spatial niche partitioning. Additionally, waterbirds are vectors of many types of organisms, both terrestrial and aquatic (40), and their propensity for moving between similar types of habitat makes them likely candidates for providing directed dispersal for hitchhikers (41–43).

A number of examples have also emerged where animal vectors respond differentially to infection by pathogens (reviewed in 44, 45). One of the best-documented examples is in malaria parasites vectored by mosquitoes, where foraging mosquitoes are more attracted to malaria-infected hosts (46–49; but see 50). There are many other disease vectors that either prefer or avoid feeding on infected hosts (51–57). Vectors preferring infected or uninfected hosts can affect host–pathogen dynamics in these systems (58–64), and consequently the stability of competitive interactions between pathogens. Given the pervasiveness of interactions between infection and microbiota (65–67), this form of dispersal may also have community-wide effects on human-associated microbes.

Another type of microbial association with evidence for animal-mediated, non-random dispersal is between nectar-associated microbes and the pollinators that disperse them from flower to flower. In the sticky monkeyflower (*Diplacus aurantiacus*), a hummingbird-pollinated shrub native to California and Oregon, the nectar of individual flowers are typically either yeast- or bacteria-dominated (68). These local alternative states are likely shaped by priority effects: early-arriving yeast (e.g., *Metschnikowia reukaufii*, *M. gruesii*, *M. rancensis*) can rapidly reduce amino acid concentrations and suppress bacteria (e.g., *Acinetobacter nectaris*, *Neokomagataea* sp.), while early-arriving bacteria reduce nectar pH and suppress yeast (68, 69). Dispersal abilities differ between yeast and bacteria, with yeast better at dispersing via pollinators, and bacteria better at dispersing in the absence of pollinators (70, 71). Belisle et al. (72) showed that plant location and nearby flower density best explained yeast colonization of sticky monkeyflower flowers, suggesting spatially non-random foraging by pollinators as a driver of yeast-dominated communities. More-recent work has shown that yeast-dominated flowers are especially attractive to pollinators, while those dominated by bacteria are repellent (73–76). In summary, nectar microbe communities of the sticky monkeyflower have local priority effects, among-species differences in dispersal, and pollinator-mediated dispersal that is affected by local communities.

Motivated by these empirical findings, here we use a mathematical model to explore how animal-mediated, non-random dispersal can stabilize regional coexistence in a system with local alternative states (Figure 1). This question is central to the understanding of species coexistence, yet few studies have addressed it. In our model, pollinators carry yeast and bacteria between flowers and leave bacteria-dominated flowers more readily than other flowers (the “microbe–pollinator feedback” hereafter). Plants are coupled only through a shared regional pool of pollinators, so we can compare competitive outcomes within plants and across the region, with and without the feedback. To test for regional coexistence, we asked whether each microbe can increase when rare in a metacommunity dominated by the other, a key requirement for stable coexistence (77–79).

**Figure 1:**
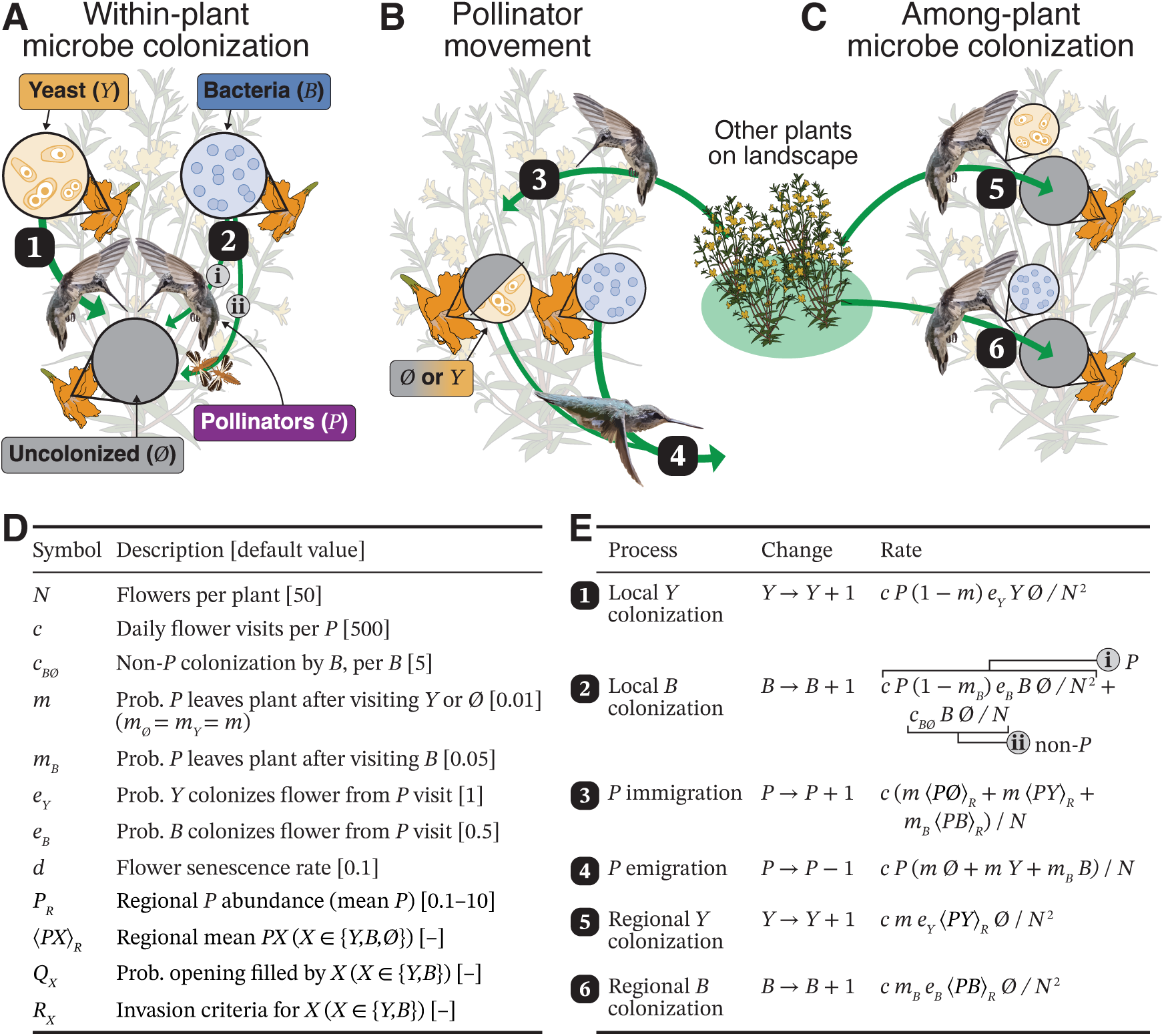
Model structure with transitions for one plant with *N* flowers. (A) The plant has uncolonized (gray, ∅Ø), yeast-dominated (yellow, *Y*), and bacteria-dominated (blue, *B*) flowers, with ∅Ø = *N* − *Y* − *B*. Flowers only transition from yeast- or bacteria-dominated to uncolonized via flower senescence followed by rapid replacement (not shown; rates are *dY* and *dB*, respectively). Yeast and bacteria both colonize uncolonized flowers on a plant via pollinator visits. Yeast colonize flowers more effectively via pollinators, but only bacteria can also colonize without pollinators. (B) Pollinators can immigrate from the surrounding landscape of plants, but are more likely to leave as bacteria-dominated flower abundance increases. (C) Incoming pollinators potentially bring microbes with them to colonize flowers on the focal plant. (D) Model parameters (with default values) and derived quantities. (E) Processes for one plant with associated state changes and rates. Rows give the rate of each process; regional colonization (5, 6) happens when an arriving pollinator colonizes a flower, so it is part of pollinator immigration (3), not additional arrivals. Boxes with numbers and lowercase Roman numerals match those in panels A–C. See SI Appendix, Table S1 for the corresponding transitions. (D,E) Rates are per day. Hummingbird images are by Rick Morris, and other illustrations are by Dr. Esther Nell. Both were used with permission.

## Modeling Framework

We consider dynamics at three increasing scales: the within-flower scale, the within-plant scale, and the among-plant field scale. At the smallest and fastest within-flower scale, populations of bacteria and yeast compete for resources and alter the pH. We do not model this scale explicitly, but based on previous work showing priority effects (68), we coarse-grain the state as either uncolonized, yeast-dominated, or bacteria-dominated. At the intermediate within-plant scale (henceforth the *local scale*), we model the dynamics of flowers and pollinators using deterministic and stochastic patch-occupancy models (80). Finally, at the largest scale (henceforth the *regional scale*), we model the dynamics of a large number of plants coupled by pollinator-driven dispersal using structured metacommunity models (20). The models at the local and regional scales are closely related and are driven by the same processes that we describe below.

At the local plant scale, we use patch-occupancy models where each patch (flower) can only be occupied by a single species. Patch-occupancy models assume that local interactions occur faster than extinction–colonization dynamics (8, 12). In our empirical system, competitive exclusion can occur in less than two days (68), and although colonization timing is unclear, flower senescence typically occurs after 7–10 days (81). We model the numbers of yeast-dominated (*Y*), bacteria-dominated (*B*), and uncolonized (∅Ø = *N* − *Y* − *B*) flowers on a plant with *N* flowers, together with the number of pollinators on that plant (*P*). We kept flower numbers constant across all plants—despite the obvious inter-plant variability in flower numbers and the effects it would have on pollinator activity—to focus on microbial competition and its interactive effects on dispersal via pollinator visits.

Each pollinator visits flowers at rate *c* (Figure 1). After visiting a flower in state *X* (*X* ∈ {∅Ø, *Y*, *B*}), the pollinator leaves the plant with probability *m_X_* and otherwise moves to another flower on the same plant. We set *m*_∅Ø_ = *m_Y_* = *m* and *m_B_* ≥ *m*: pollinators leave bacteria-dominated flowers more readily because bacteria make nectar acidic and less palatable (68, 73, 74). The ratio *m_B_*/*m* therefore sets the strength of the microbe–pollinator feedback, and there is no feedback when *m_B_* = *m*. A pollinator arriving at an uncolonized flower from a yeast- or bacteria-dominated flower establishes that microbe with probability *e_Y_* or *e_B_*. Bacteria (but not yeast) also colonize uncolonized flowers without pollinators, via thrips and other non-pollinating insects (71), at rate *c_B_*_∅Ø_ per bacteria-dominated flower. Flowers senesce at rate *d* and are immediately replaced by uncolonized flowers to keep *N* constant.

Pollinators that leave a plant join a regional pool and arrive at other plants at random. The regional pollinator abundance *P_R_* is the mean number of pollinators per plant. Because the probability of a pollinator leaving a plant depends on its microbial community, we have to describe the pool by ⟨*P*∅Ø⟩*_R_*, ⟨*PY*⟩*_R_*, and ⟨*PB*⟩*_R_*, the means across plants of *P*∅Ø, *PY*, and *PB*, where ⟨*P*∅Ø⟩*_R_* + ⟨*PY*⟩*_R_* + ⟨*PB*⟩*_R_* = *NP_R_*. Thus pollinators that last visited an *X* flower on another plant arrive at the focal plant at rate *c m_X_* ⟨*PX*⟩*_R_*/*N*. The resulting transitions are in Figure 1E.

Following (20), we embed these processes in a family of related models. We start by considering a deterministic patch-dynamics model of flowers on a single focal plant, first in isolation, then open to dispersal with the region. We then introduce demographic stochasticity due to the small number of flowers and pollinators per plant in a related continuous-time Markov process. Finally, we consider a closed regional metacommunity of a large number of plants coupled by dispersal, using the structured metacommunity approach of (20) to find regional equilibria and calculate invasion criteria to determine the outcome of competition (see *Methods* for details).

### Reduced models

Pollinators visit flowers much faster than flowers senesce (*c* = 500 versus *d* = 0.1 per day), so uncolonized flowers are refilled almost immediately. To reduce the dimensionality of the system to aid understanding, we use a reduced model with no uncolonized flowers (∅Ø = 0, so *B* = *N* − *Y*), in which a senescing yeast-dominated flower is replaced by a bacteria-dominated flower or vice versa. This assumption is common to lottery models (82) and Hubbell’s netural model (83) and does not meaningfully affect our results (SI Appendix, Numerical accuracy). See *Methods* for more details on this simplifying assumption.

## Results

### Local-scale within-plant dynamics

In the simplest case of a single isolated plant with pollinator numbers fixed and no microbe immigration (*m_B_* = *m* = 0), priority effects at the flower scale up to competitive exclusion at the plant scale, as microbes compete for empty flowers (80). The winner of competition depends on the number of pollinators. There is a critical density of pollinators *P*_crit_. Because bacteria can colonize flowers by thrips in the absence of pollinators, they exclude yeast when *P* < *P*_crit_.

However yeast benefit from pollinators more, so they exclude bacteria when *P* > *P*_crit_ (Figure 2A).

**Figure 2:**
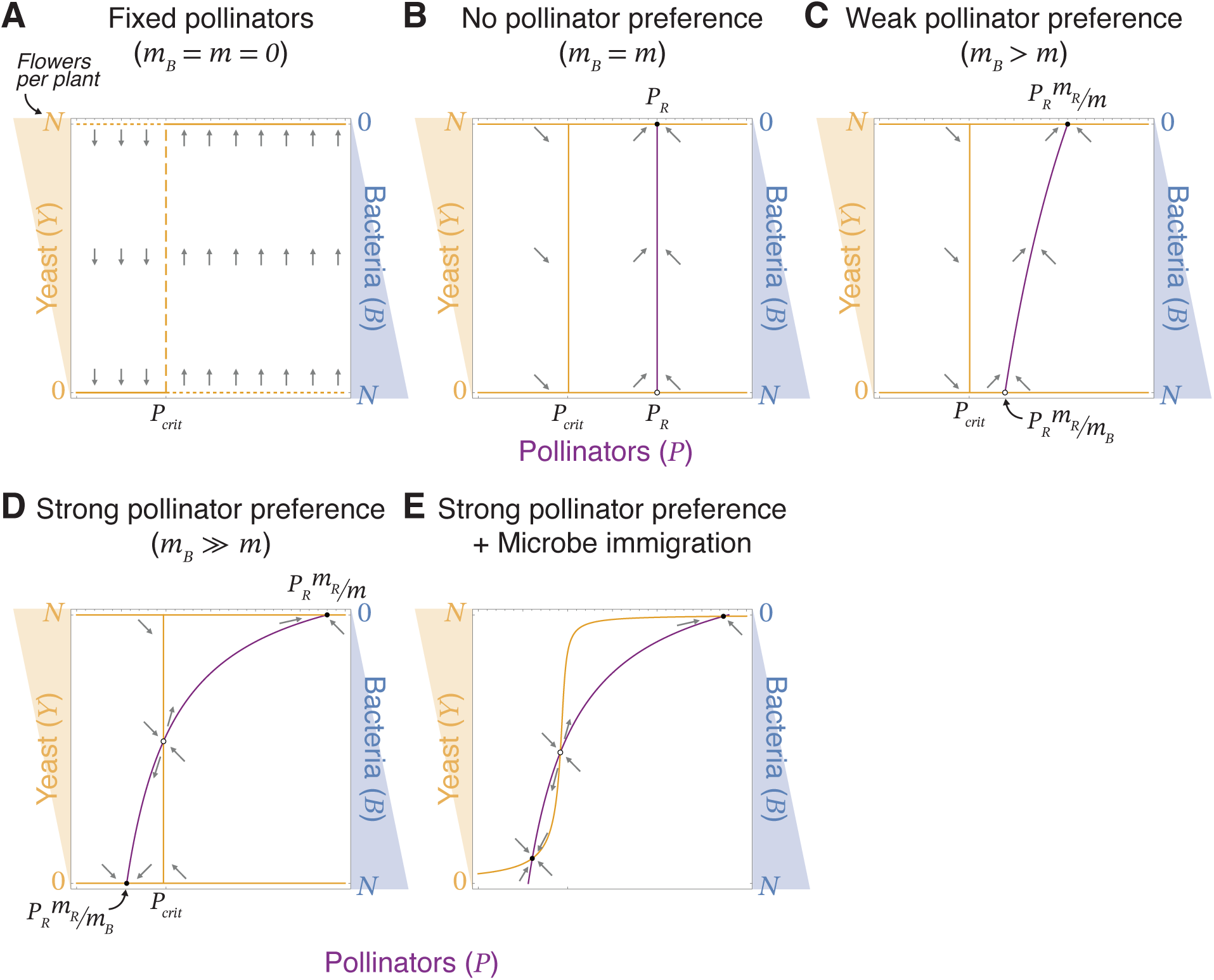
Single-plant deterministic model (equation 3). Shown are phase-plane diagrams of pollinators (*P*, x-axis) versus yeast-dominated flowers (*Y*, left y-axis), with *B* = *N* − *Y* on the right y-axis. Orange and purple curves are yeast and pollinator isoclines; filled and open points are stable and unstable equilibria. (A) With pollinators fixed (*m_B_* = *m* = 0), bacteria win if *P* < *P*_crit_ and yeast win if *P* > *P*_crit_. Note that in this panel, *P* is a fixed parameter, so it shows the direction of change in *Y* as a one-dimensional flow drawn in the plane. (B–D) Pollinators move between the plant and the regional pool but do not carry microbes. In these panels, *P* is a dynamic variable. (B) Without a pollinator preference (*m_B_* = *m*), local pollinator abundance equals the regional abundance, *P_R_*, so bacteria win if *P_R_* < *P*_crit_ and yeast win if *P_R_* > *P*_crit_. (C) With a weak pollinator preference for yeast-dominated flowers (*m_B_* > *m*), there is a single stable equilibrium. (D) With a strong pollinator preference (*m_B_* ≫ *m*), there are two alternative stable equilibria: all yeast with *P* > *P*_crit_, or all bacteria with *P* < *P*_crit_. (E) The alternative stable equilibria remain when pollinators also carry microbes between plants, resulting in mass effects. (A–E) *m_R_* = (*m*⟨*PY*⟩*_R_* + *m_B_* ⟨*PB*⟩*_R_*)/(*NP_R_*) is the composition-weighted emigration probability of the regional pool. (B–E) *P_R_* = 2 and ratio of ⟨*PY*⟩*_R_* to ⟨*PB*⟩*_R_* is 9, with *m_B_* = 0.01 (B), 0.015 (C), or 0.05 (D, E).

Letting pollinators move between the focal plant and the regional pool, but neglecting microbe immigration, makes the equilibrium pollinator abundance *P* potentially depend on plant composition:

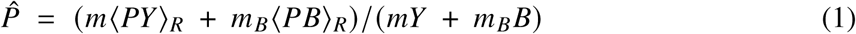

When the pollinator does not respond to microbes (*m_B_* = *m*), *P*^^^ = *P_R_* regardless of composition, resulting in the same outcome of competitive exclusion as described above, with *P_R_* in place of *P* (Figure 2B). When pollinators respond to microbes (*m_B_* > *m*), their local abundance is higher than their regional abundance on yeast-dominated plants (*P*^^^ = *P_R_m_R_*/*m*) but lower than their regional abundance on bacteria-dominated plants (*P*^^^ = *P_R_m_R_*/*m_B_*), where *m_R_* = (*m*⟨*PY*⟩*_R_* + *m_B_* ⟨*PB*⟩*_R_*)/(*NP_R_*) is the mean emigration probability of the regional pool. A weak preference for yeast-dominated flowers leaves a single stable equilibrium (Figure 2C).

However when the preference is strong enough that *P_R_m_R_*/*m_B_* < *P*_crit_ < *P_R_m_R_*/*m*, the plant has two alternative stable states: yeast-dominated (*Y* = *N*) with many pollinators, or bacteria-dominated (*Y* = 0) with few (Figure 2D). These alternative stable states persist when pollinators also carry microbes into the plant (Figure 2E).

With demographic stochasticity due to finite flowers and pollinator abundances, these two stable states become the two modes of a bimodal stationary probability distribution (Figure 3C). Plants started near either state remain there for 1000 days (Figure 3A,B), although over longer periods they switch between states. Thus a sufficiently strong microbe–pollinator feedback can convert plant-scale competitive exclusion into plant-scale priority effects.

**Figure 3:**
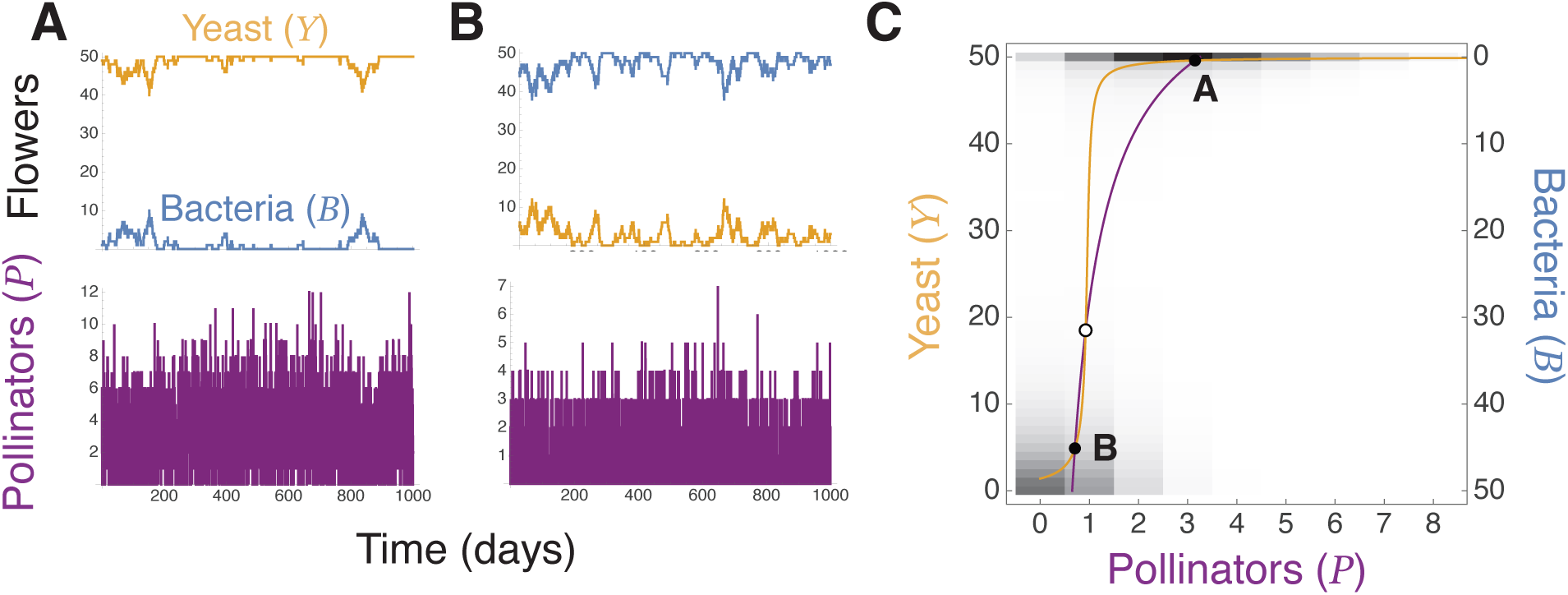
Alternative stable states in the deterministic model become a bimodal stationary distribution in the stochastic model (vacancy closure). (A,B) Simulations started near each stable state (A: *Y* = 49, *P* = 3; B: *Y* = 5, *P* = 1) remain near it for 1000 days, although plants eventually switch between states. (C) Deterministic phase plane, as in Figure 2, over the stationary distribution of (*Y*, *P*) (grayscale). *P_R_* = 1.8, with ⟨*PY*⟩*_R_* = 71.87 and ⟨*PB*⟩*_R_* = 18.13 at the metacommunity equilibrium.

### Regional-scale among-plant metacommunity

In the closed metacommunity, bacteria exclude yeast at low regional pollinator abundance (*P_R_* < 0.73), yeast exclude bacteria at high abundance (*P_R_* > 2.90), and the two stably coexist in between (Figures 4, 5). Within the coexistence range, each plant is dominated by one species most of the time, making the metacommunity a mosaic of yeast-dominated plants with many pollinators and bacteria-dominated plants with few (Figure 4B–D). As *P_R_* increases, the proportion of yeast-dominated plants increases from 0 to 1 (Figure 5D), while mean pollinator numbers stay near 3 on yeast-dominated plants and near 0.7 on bacteria-dominated plants (Figure 5E). These values lie above and below *P*_crit_ = 1, respectively, so each type of plant keeps the pollinator number at which its dominant microbe wins locally. Increasing *P_R_* therefore changes the proportions of the two plant types more than it changes the distribution of microbes on the plants themselves.

**Figure 4:**
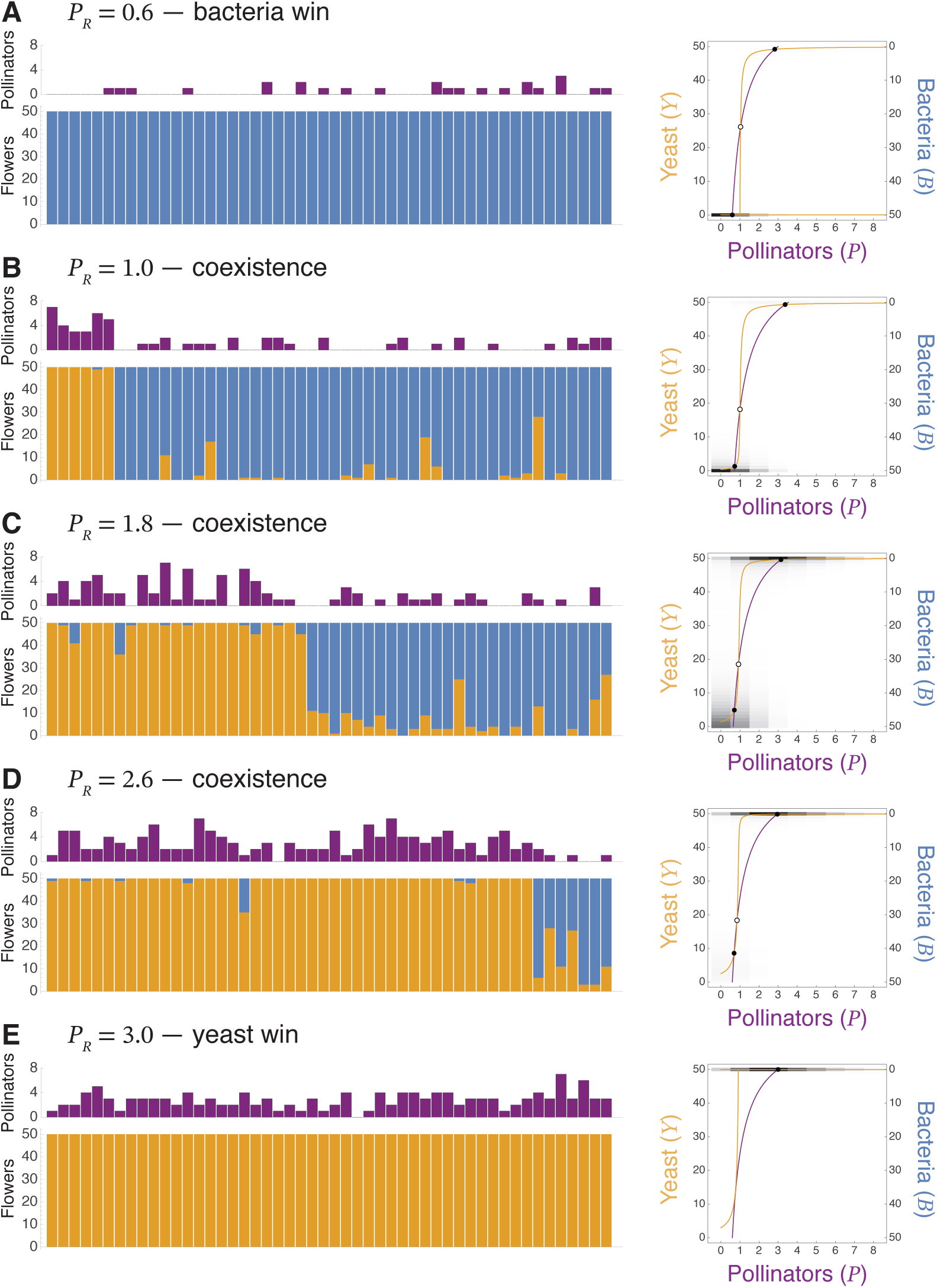
Equilibria of the closed metacommunity. (A) Bacteria exclude yeast at low regional pollinator abundance, *P_R_*. (B–D) Yeast and bacteria coexist at intermediate *P_R_*. (E) Yeast exclude bacteria at high *P_R_*. Left panels: 50 plants drawn at random from the stationary distribution (top, pollinators; bottom, yeast- and bacteria-dominated flowers). Right panels: stationary distribution of one plant (grayscale) with the deterministic phase plane.

**Figure 5:**
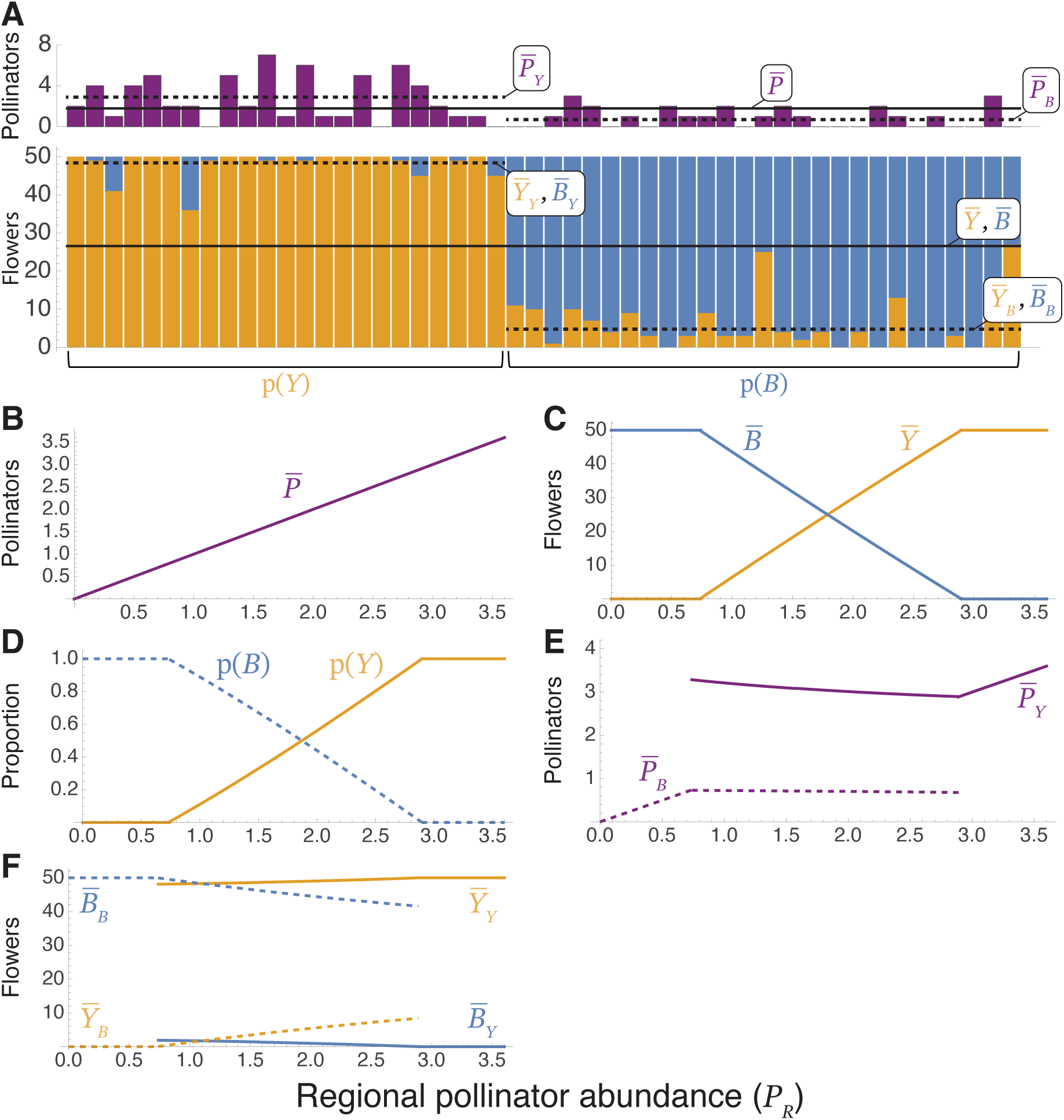
Community structure along a gradient of regional pollinator abundance, *P_R_*, in the closed metacommunity. (A) Summary statistics, illustrated with 50 plants drawn from the stationary distribution: means across all plants (*P̅*, *Y̅*, *B̅*), proportions of yeast- and bacteria-dominated plants (p(*Y*), p(*B*)), and means within yeast- and bacteria-dominated plants (subscripts *Y* and *B*). (B) Mean pollinators per plant. (C) Mean yeast- and bacteria-dominated flowers per plant. (D) Proportions of yeast- and bacteria-dominated plants. (E) Mean pollinators on yeast- and bacteria-dominated plants. (F) Mean yeast- and bacteria-dominated flowers on yeast- and bacteria-dominated plants. Yeast and bacteria coexist for 0.73 < *P_R_* < 2.90.

The range of *P_R_* allowing coexistence widens as pollinators respond more strongly to bacteria (Figures 6, S1); without a pollinator response (*m_B_* = *m* = 0.01), both species could mutually invade only for the narrow range of 1.46 < *P_R_* < 1.57. The coexistence region lies largely inside the region where a plant has alternative stable states in the deterministic model without microbe immigration, *m_B_*/*m* > max(*P_R_*/*P*_crit_, *P*_crit_/*P_R_*) (dashed curves in Figure 6). However, plant-scale alternative stable states are not sufficient for coexistence, since much of that region leads to exclusion, nor strictly necessary, since coexistence extends outside it near *m_B_* = *m*.

**Figure 6:**
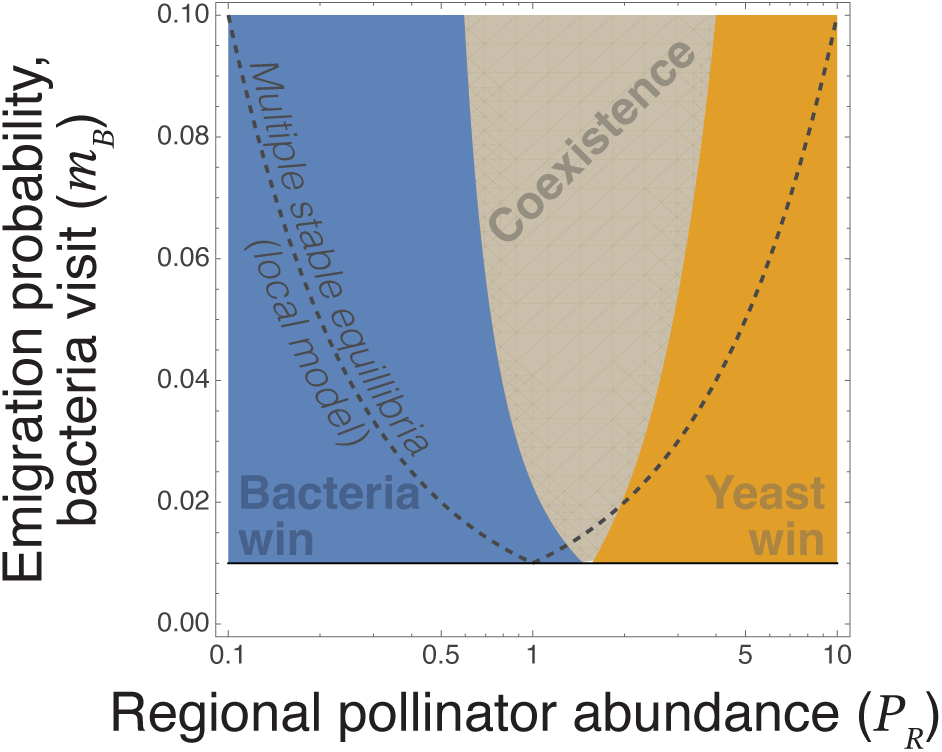
Outcome of competition in relation to regional pollinator abundance, *P_R_*, and the emigration probability from bacteria-dominated flowers, *m_B_*, which sets the strength of the microbe–pollinator feedback, for *m_B_* ≤ 0.1. Fill indicates outcomes, which are from invasion criteria (Methods). The horizontal line at the bottom of each pane is *m_B_* = *m* = 0.01 (no pollinator preference). Dashed curves bound the region where a plant has alternative stable equilibria in the deterministic model without microbe immigration, *m_B_*/*m* > max(*P_R_*/*P*_crit_, *P*_crit_/*P_R_*). Alternative stable equilibria are neither sufficient nor strictly necessary for regional coexistence, but the coexistence region lies largely within the region of multiple stable equilibria. Figure S1 shows these outcomes for *m_B_* ≤ 1.

## Discussion

Overall, our results indicate that non-random dispersal can facilitate regional coexistence, not just in spite of local priority effects, but because of them. Specifically, our modeling work suggests the following dispersal–community hypothesis: In competing nectar microbes, individual flowers can be dominated by either bacteria or yeast, with the early-arriving species excluding the other. When pollinator-mediated dispersal is affected by the communities of nectar microbes on a given plant, this feedback causes plant-level priority effects as well (Figure 2). However, in metacommunities where plants share a regional pool of pollinators, the microbe–pollinator feedback reinforces the spatial mosaic of plants with alternative stable states (Figures 4, 5). This spatial heterogeneity generates a spatial niche partitioning (84), where each species can proliferate in locations that are advantageous to them (low- and high-pollination plants for bacteria and yeast, respectively). A stronger microbe–pollinator feedback increases the difference in pollinator visitation between common and rare species in the direction that benefits whichever is rare. This feedback makes landscape-wide rarity increasingly beneficial, strengthening spatial niche partitioning and the ability of each species to increase in abundance when rare. The microbe–pollinator feedback thereby promotes stable coexistence (Figure 6). In other words, destabilizing positive feedback at the flower and plant scales yield stabilizing negative feedback at the landscape scale, causing coexistence at this scale even when exclusion is what we see at smaller scales.

Although we used global dispersal in our model, a spatially explicit model with continuous among-plant microbial dispersal would add a layer of realism and help determine how robust our results are to patterns of connectivity among plants. Placing plants randomly or uniformly on a two-dimensional landscape and having dispersal decay with distance would make coexistence difficult because it would cause regional abundance to dictate among-plant dispersal, causing regional positive density-dependence (19). On the other hand, incorporating hummingbird behavior, such as territoriality and foraging via “trap-lining” (85, 86), could yield interesting forms of spatial connectivity among plants that could further bolster the coexistence mechanisms shown here. Additionally, letting the number of flowers vary among plants would allow for tests of the interactive effects of how pollinators respond not only to microbial communities, but also to flower densities.

In our metacommunity model, the regional pollinator pool is closed: regional pollinator abundance, *P_R_*, is fixed, so pollinators that leave bacteria-dominated plants are redistributed among other plants. When among-plant variation in bacterial abundance is combined with a microbe–pollinator feedback, plants with lower abundances of bacteria pulled pollinator visits away from plants with more bacteria-dominated flowers. This disparity in pollinator visits coincided with the conditions that reduced competition for each species. Because pollinator numbers on each plant depended on its own microbial composition relative to that of the regional pool, a single plant dominated by an invading (i.e., rare) microbial competitor would experience drastically different pollinator visitations compared to resident-dominated plants. This caused each species to increase when rare. Visits by animal vectors can be limiting for many other dispersal-limited organisms, including plants with animal-dispersed seeds (87, 88), vector-borne pathogens (89), and organisms living in ephemeral habitats (90). When this limited dispersal is also directed and creates opportunities for reduced competition across space, it could promote coexistence in these systems.

In the nectar microbe system modeled here, the differences among community states in their desirability for animal vectors is mediated by the different effects of competitors on the abiotic conditions of each habitat patch. Specifically, bacteria make nectar acidic and likely foul-tasting to pollinators, while yeast does not (68, 73, 74). This contrast parallels pathogens that alter volatile chemicals produced by their host to increase visitations by vectors (46–49, 52–56). Competitive outcomes among pathogens at an epidemiological scale could be altered if one or more species can alter vector behavior to facilitate their own movement to new hosts. For example, novel host odors that are preferential to vectors could cause negative frequency-dependence in competition for vector dispersal, stabilizing coexistence. Another way pathogen manipulation of vectors could contribute to among-pathogen coexistence is through a difference among pathogens in their reliance on one or more vector species, combined with pathogens inducing odor profiles that preferentially attract those vectors that best disperse them. This difference would cause spatial niche partitioning similar to that studied here. Two key properties that may indicate the presence of this type of spatial niche partitioning are among-species differences in animal-mediated dispersal and reduced competition in environmentally favorable locations (35).

The applicability of our results to real communities in the field may be limited by a number of factors. Changes in animal-mediated dispersal must alter the outcome of competition for our mechanism to work. In field settings, however, effects of animal-mediated dispersal may be overwhelmed by the many other properties governing competitive outcomes. Field experiments where animal vectors are excluded (as in 70) can demonstrate whether changes in vector-mediated dispersal alter competition in the field. Additionally, directed dispersal is required for the mechanism we studied here, but in more complex communities, one or more interacting species dispersing indiscriminately could negate any effects of non-random dispersal on coexistence.

Field experiments where treatments alter starting community compositions (as in 75) could test for how robust the predictions are to non-focal animal vectors that might not be responding to community conditions.

Our study has shown how non-random dispersal mediated by animal vectors can promote stable regional coexistence despite local priority effects. Many organisms rely on animals for dispersal, and evidence suggests that these animal vectors often disperse in response to the environmental changes the dispersed organisms bring about. Our modeling suggests that this form of dispersal–community feedback can change fundamental properties of ecological communities.

## Methods

### Reduced model

To eliminate uncolonized flowers ∅Ø from the model, we compute the rates at which a single uncolonized flower would be colonized by yeast (*a_Y_*) and bacteria (*a_B_*):

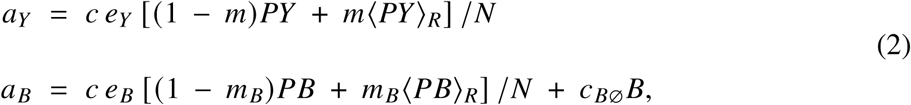

The resulting deterministic reduced model is

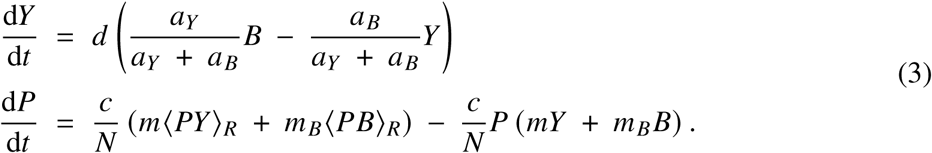

Calculating the transition rates in the stochastic reduced model is more complicated. When the state of a plant is *Y*, *Y* increases by one at rate *d B Q_Y_* (*Y*), when a bacteria-dominated flower senesces and the opening is filled by yeast, and decreases by one at rate *d Y Q_B_* (*Y*). *Q_Y_* and *Q_B_* are the probabilities that the opening is eventually filled by yeast or bacteria. Because pollinators can arrive and leave while the flower is open, we must solve for *Q_Y_* (*Y*) and *Q_B_* (*Y*) to use in our stochastic model.

Our assumption of no uncolonized flowers requires the timescale of recolonization to be much less than the timescale of flower senescence. However, in the stochastic model, a third timescale must be considered: the timescale of pollinator dispersal. We first describe two limiting cases. In one limiting case, if the timescale of pollinator dispersal is large relative to the timescale of recolonization, then the number of pollinators on a plant can be treated as constant while the flower is empty. In another limiting case, if the timescale of pollinator dispersal is small relative to the timescale of recolonization, then the pollinators will fluctuate and approach a Poisson distribution governed by the balance of immigration and emigration.

When the timescales of recolonization and pollinator dispersal are not cleanly separated, we can compute the probabilities *Q_Y_* (*Y*) and *Q_B_* (*Y*) with a Markov chain for the open flower: plant composition is held fixed, *P* changes by pollinator immigration and emigration, and the opening is filled at the colonization rates in Figure 1E. The filling probabilities from each starting *P* solve a tridiagonal linear system (SI Appendix), and we averaged them over the pollinator count at the moment of senescence, taken as *P* | *Y* ∼ Poisson(*P*^^^(*Y*)). We call this the vacancy closure. The stationary distribution of *Y* follows from detailed balance of this birth–death process, and we reconstructed the joint distribution of (*Y*, *P*) with *P* | *Y* ∼ Poisson(*P*^^^(*Y*)). We use this vacancy closure in our results, which closely match the full model with uncolonized flowers, ∅Ø (SI Appendix, Numerical accuracy).

### Stochastic patch-occupancy model

We use two methods to analyze the stochastic patch-occupancy model. To numerically calculate the stationary probability distribution as in Figure 3C, we construct a state-transition matrix based on the transition rates, then find the eigenvector associated with its dominant zero eigenvalue. To simulate time series, we simulated *Y* and *P* with the pollinator immigration and emigration rates using the Gillespie algorithm (91).

### Closed metacommunity

We model many (effectively infinite) plants coupled only through the regional pollinator pool, adapting the closed-metacommunity method of Lerch et al. (20). The key idea is that the regional abundances ⟨*PY*⟩*_R_* and ⟨*PB*⟩*_R_* must equal to their mean local abundances ⟨*PY*⟩ and ⟨*PB*⟩, averaged over the stationary distribution of the open stochastic model to be consistent. The regional pollinator abundance *P_R_* is fixed, and the composition of the pool is set by the plants: ⟨*PY*⟩*_R_* = ⟨*PY*⟩ and ⟨*PB*⟩*_R_* = ⟨*PB*⟩ = *NP_R_* − ⟨*PY*⟩*_R_*, where expectations are over the stationary distribution of one plant given the pool, which for many plants is also the distribution of plant states across the metacommunity (20). We solved ⟨*PY*⟩*_R_* = ⟨*PY*⟩ numerically using Newton’s method, and traced the solution across *P_R_* in steps of 0.01 from *P_R_* = 1. Plants in a metacommunity at equilibrium differ because the stationary distribution of a plant can be bimodal. To summarize the metacommunity in Figure 5, we split the stationary distribution of (*Y*, *P*) between its modes along the watershed between them, which gives the proportions of yeast- and bacteria-dominated plants, *p*(*Y*) and *p*(*B*), and the means within each.

To determine the outcome of competition at the regional scale, we use invasion analysis (92). To calculate invasion criteria, we solve for a monoculture of one microbe, then introduce a small seed of the other. Yeast can invade a bacteria-only metacommunity if a small pool of yeast-carrying pollinators, ⟨*PY*⟩*_R_* = *ε*, produces a larger one: *R_Y_* = ⟨*PY*⟩/*ε* > 1 (20), evaluated with ⟨*PB*⟩*_R_* = *NP_R_* − *ε*. Likewise, bacteria can invade a yeast-only metacommunity if *R_B_* = ⟨*PB*⟩/*ε* > 1 with ⟨*PY*⟩*_R_* = *NP_R_* − *ε*. We used *ε* = 10^−5^ and classified outcomes as bacteria win (*R_Y_* < 1 < *R_B_*), coexistence by mutual invasibility (*R_Y_* > 1 and *R_B_* > 1), or yeast win (*R_B_* < 1 < *R_Y_*). With *m_B_* ≥ *m*, there were no parameter values at which neither species could invade, the regional founder control of ref. (20); this outcome requires *m_B_* < 0.0091 (scan over 0.1 ≤ *P_R_* ≤ 10). The location of *R* = 1 does not depend on *ε* or *P*_max_, but the magnitude of *R* does, so we report only whether each species can invade (SI Appendix, Table S2).

### Parameter values

Unless stated otherwise, we used *N* = 50 flowers per plant, *c* = 500 day^−1^, *d* = 0.1 day^−1^, *m* = 0.01, *m_B_* = 0.05, *e_Y_* = 1, *e_B_* = 0.5, and *c_B_*_∅Ø_ = 5 day^−1^, for which *P*_crit_ = 1. These values were based on our empirical knowledge of the system (T. Fukami, unpublished results). For numerical solutions we truncated the pollinator count at *P*_max_ = 12.

### Software

For model simulations and analyses, we used the EcoEvo package v1.7.2 (https://github.com/cklausme/EcoEvo) on Wolfram v15.0.1 (93). The figures were made in Wolfram. For open access, we also wrote an R (94) package and scripts that reproduce every computed result; simulations and random draws (Figures 3A,B and 4) match in distribution only.

## Acknowledgments

This material is based upon work supported by the National Science Foundation Postdoctoral Research Fellowships in Biology Program under Grant No. 2209354. T.F. acknowledges support from the National Science Foundation (DEB-1737758 and OIE-2618565). C.A.K. acknowledges support from the National Science Foundation (NSF EF-2124800).

## References

[1] Levin, S. A. (1992) The problem of pattern and scale in ecology. Ecology 73, 1943–1967.

[2] Chesson, P. (1996) in Frontiers of Population Ecology, eds. Floyd, R. B, Sheppard, A. W, & De Barro, P. J. (CSIRO Publishing, Melbourne, Australia), pp. 353–368.

[3] Marino, P. C. (1988) Coexistence on divided habitats: Mosses in the family Splachnaceae. Annales Zoologici Fennici 25, 89–98.

[4] Levin, S. A. (2000) Multiple scales and the maintenance of biodiversity. Ecosystems 3, 498–506.

[5] Hassell, M. P, Comins, H. N, & May, R. M. (1994) Species coexistence and self-organizing spatial dynamics. Nature 370, 290–292.

[6] Calcagno, V, Mouquet, N, Jarne, P, & David, P. (2006) Coexistence in a metacommunity: The competition-colonization trade-off is not dead. Ecology Letters 9, 897–907.

[7] Levins, R & Culver, D. (1971) Regional coexistence of species and competition between rare species. Proceedings of the National Academy of Sciences 68, 1246–1248.

[8] Tilman, D. (1994) Competition and biodiversity in spatially structured habitats. Ecology 75, 2–16.

[9] Amarasekare, P. (2004) Spatial variation and density-dependent dispersal in competitive coexistence. Proceedings of the Royal Society of London. Series B: Biological Sciences 271, 1497–1506.

[10] Levin, S. A. (1974) Dispersion and population interactions. The American Naturalist 108, 207–228.

[11] Horn, H. S & MacArthur, R. H. (1972) Competition among fugitive species in a harlequin environment. Ecology 53, 749–752.

[12] Amarasekare, P & Nisbet, R. M. (2001) Spatial heterogeneity, source-sink dynamics, and the local coexistence of competing species. The American Naturalist 158, 572–584.

[13] Chase, J. M. (2003) Community assembly: When should history matter? Oecologia 136, 489–498.

[14] Fukami, T. (2015) Historical contingency in community assembly: Integrating niches, species pools, and priority effects. Annual Review of Ecology, Evolution, and Systematics 46, 1–23.

[15] Fukami, T, Mordecai, E. A, & Ostling, A. (2016) A framework for priority effects. Journal of Vegetation Science 27, 655–657.

[16] Letten, A. D, Ke, P.-J, & Fukami, T. (2017) Linking modern coexistence theory and contemporary niche theory. Ecological Monographs 87, 161–177.

[17] Mordecai, E. A. (2011) Pathogen impacts on plant communities: Unifying theory, concepts, and empirical work. Ecological Monographs 81, 429–441.

[18] Ke, P.-J & Letten, A. D. (2018) Coexistence theory and the frequency-dependence of priority effects. Nature Ecology and Evolution 2, 1691–1695.

[19] Shurin, J. B, Amarasekare, P, Chase, J. M, Holt, R. D, Hoopes, M. F, & Leibold, M. A. (2004) Alternative stable states and regional community structure. Journal of Theoretical Biology 227, 359–368.

[20] Lerch, B. A, Rudrapatna, A, Rabi, N, Wickman, J, Koffel, T, & Klausmeier, C. A. (2023) Connecting local and regional scales with stochastic metacommunity models: Competition, ecological drift, and dispersal. Ecological Monographs 93, e1591.

[21] Wittmann, M. J & Fukami, T. (2018) Eco-evolutionary buffering: Rapid evolution facilitates regional species coexistence despite local priority effects. The American Naturalist 191, E171–E184.

[22] Nell, L. A, Weng, Y.-M, Phillips, J. S, Botsch, J. C, Book, K. R, Einarsson, A, Ives, A. R, & Schoville, S. D. (2024) Shared features underlying compact genomes and extreme habitat use in chironomid midges. Genome Biology and Evolution 16, evae086.

[23] Amarasekare, P. (2007) Spatial dynamics of communities with intraguild predation: The role of dispersal strategies. The American Naturalist 170, 819–831.

[24] Amarasekare, P, Hoopes, M. F, Mouquet, N, & Holyoak, M. (2004) Mechanisms of coexistence in competitive metacommunities. The American Naturalist 164, 310–326.

[25] Amarasekare, P. (2010) Effect of non-random dispersal strategies on spatial coexistence mechanisms. Journal of Animal Ecology 79, 282–293.

[26] Lin, W.-T, Hsieh, C.-h, & Miki, T. (2013) Difference inadaptive dispersal ability can promote species coexistence in fluctuating environments. PLoS ONE 8, e55218.

[27] Zhang, H, Zhu, Y, Liu, Z, Peng, Y, Peng, W, Tong, L, Wang, J, Liu, Q, Wang, P, & Cheng, G. (2022) A volatile from the skin microbiota of flavivirus-infected hosts promotes mosquito attractiveness. Cell 185, 2510–2522.e16.

[28] Custer, G. F, Bresciani, L, & Dini-Andreote, F. (2022) Ecological and evolutionary implications of microbial dispersal. Frontiers in Microbiology 13, 855859.

[29] Baltzinger, C, Karimi, S, & Shukla, U. (2019) Plants on the move: Hitch-hiking with ungulates distributes diaspores across landscapes. Frontiers in Ecology and Evolution 7, 38.

[30] Bartlow, A. W & Agosta, S. J. (2021) Phoresy in animals: Review and synthesis of a common but understudied mode of dispersal. Biological Reviews 96, 223–246.

[31] Berkley, H. A, Kendall, B. E, Mitarai, S, & Siegel, D. A. (2010) Turbulent dispersal promotes species coexistence. Ecology Letters 13, 360–371.

[32] Ives, A. R. (1988) Covariance, coexistence and the population dynamics of two competitors using a patchy resource. Journal of Theoretical Biology 133, 345–361.

[33] Ives, A. R. (1991) Aggregation and coexistence in a carrion fly community. Ecological Monographs 61, 75–94.

[34] Ruokolainen, L & Hanski, I. (2016) Stable coexistence of ecologically identical species: Conspecific aggregation via reproductive interference. Journal of Animal Ecology 85, 638–647.

[35] Chesson, P. L. (2000) General theory of competitive coexistence in spatially-varying environments. Theoretical Population Biology 58, 211–237.

[36] Davidson, D. W & Morton, S. R. (1981) Competition for dispersal in ant-dispersed plants. Science 213, 1259–1261.

[37] Davidson, D. W & Morton, S. R. (1981) Myrmeeoehory in some plants (*F. chenopodiaceae*) of the Australian arid zone. Oecologia 50, 357–366.

[38] Handel, S. N. (1978) The competitive relationship of three woodland sedges and its bearing on the evolution of ant-dispersal of *Carex pedunculata*. Evolution 32, 151–163.

[39] Docters van Leeuwen, WM. (1954) On the biology of some Javanese *Loranthaceae* and the role birds play in their life-historie. Beaufortia 4, 103–207.

[40] Green, A. J, Lovas-Kiss, A, Reynolds, C, Sebastián-González, E, Silva, G. G, Van Leeuwen, C. H. A, & Wilkinson, D. M. (2023) Dispersal of aquatic and terrestrial organisms by waterbirds: A review of current knowledge and future priorities. Freshwater Biology 68, 173–190.

[41] Almeida, B. A, Lukacs, B. A, Lovas-Kiss, A, Reynolds, C, & Green, A. J. (2022) Functional traits drive dispersal interactions between European waterfowl and seeds. Frontiers in Plant Science 12, 795288.

[42] Kleyheeg, E, Treep, J, De Jager, M, Nolet, B. A, & Soons, M. B. (2017) Seed dispersal distributions resulting from landscape-dependent daily movement behaviour of a key vector species, *Anas platyrhynchos*. Journal of Ecology 105, 1279–1289.

[43] Martın-Velez, V, Van Leeuwen, C. H. A, Sanchez, M. I, Hortas, F, Shamoun-Baranes, J, Thaxter, C. B, Lens, L, Camphuysen, C. J, & Green, A. J. (2021) Spatial patterns of weed dispersal by wintering gulls within and beyond an agricultural landscape. Journal of Ecology 109, 1947–1958.

[44] Javed, N, Bhatti, A, & Paradkar, P. N. (2021) Advances in understanding vector behavioural traits after infection. Pathogens 10, 1376.

[45] Gandon, S. (2018) Evolution and manipulation of vector host choice. The American Naturalist 192, 23–34.

[46] De Boer, J. G, Robinson, A, Powers, S. J, Burgers, S. L. G. E, Caulfield, J. C, Birkett, M. A, Smallegange, R. C, Van Genderen, P. J. J, Bousema, T, Sauerwein, R. W, Pickett, J. A, Takken, W, & Logan, J. G. (2017) Odours of *Plasmodium falciparum*-infected participants influence mosquito-host interactions. Scientific Reports 7, 9283.

[47] De Moraes, C. M, Stanczyk, N. M, Betz, H. S, Pulido, H, Sim, D. G, Read, A. F, & Mescher, M. C. (2014) Malaria-induced changes in host odors enhance mosquito attraction. Proceedings of the National Academy of Sciences 111, 11079–11084.

[48] Lacroix, R, Mukabana, W. R, Gouagna, L. C, & Koella, J. C. (2005) Malaria infection increases attractiveness of humans to mosquitoes. PLoS Biology 3, e298.

[49] Robinson, A, Busula, A. O, Voets, M. A, Beshir, K. B, Caulfield, J. C, Powers, S. J, Verhulst, N. O, Winskill, P, Muwanguzi, J, Birkett, M. A, Smallegange, R. C, Masiga, D. K, Mukabana, W. R, Sauerwein, R. W, Sutherland, C. J, Bousema, T, Pickett, J. A, Takken, W, Logan, J. G, & De Boer, J. G. (2018) *Plasmodium*-associated changes in human odor attract mosquitoes. Proceedings of the National Academy of Sciences 115, E4209–E4218.

[50] Vantaux, A, De Sales Hien, D. F, Yameogo, B, Dabire, K. R, Thomas, F, Cohuet, A, & Lefevre, T. (2015) Host-seeking behaviors of mosquitoes experimentally infected with sympatric field isolates of the human malaria parasite *Plasmodium falciparum*: No evidence for host manipulation. Frontiers in Ecology and Evolution 3, 86.

[51] Cellini, A, Giacomuzzi, V, Donati, I, Farneti, B, Rodriguez-Estrada, M. T, Savioli, S, Angeli, S, & Spinelli, F. (2019) Pathogen-induced changes in floral scent may increase honeybee-mediated dispersal of *Erwinia amylovora*. The ISME Journal 13, 847–859.

[52] Van Duijvendijk, G, Van Andel, W, Fonville, M, Gort, G, Hovius, J. W, Sprong, H, & Takken, W. (2016) A *Borrelia afzelii* infection increases larval tick burden on *Myodes glareolus* (Rodentia: Cricetidae) and nymphal body weight of *Ixodes ricinus* (Acari: Ixodidae). Journal of Medical Entomology p. tjw157.

[53] Nevatte, T. M, Ward, R. D, Sedda, L, & Hamilton, J. G. C. (2017) After infection with *Leishmania infantum*, golden hamsters (*Mesocricetus auratus*) become more attractive to female sand flies (*Lutzomyia longipalpis*). Scientific Reports 7, 6104.

[54] Baylis, M & Nambiro, C. O. (1993) The effect of cattle infection by *Trypanosoma congolense* on the attraction, and feeding success, of the tsetse fly *Glossina pallidipes*. Parasitology 106, 357–361.

[55] Coleman, R. E & Edman, J. D. (1988) Feeding-site selection of *Lutzomyia longipalpis* (Diptera: Psychodidae) on mice infected with *Leishmania mexicana amazonensis*. Journal of Medical Entomology 25, 229–233.

[56] O’Shea, B, Rebollar-Tellez, E, Ward, R, Hamilton, J, El Naiem, D, & Polwart, A. (2002) Enhanced sandfly attraction to *Leishmania*-infected hosts. Transactions of the Royal Society of Tropical Medicine and Hygiene 96, 117–118.

[57] Verhulst, N. O, Qiu, Y. T, Beijleveld, H, Maliepaard, C, Knights, D, Schulz, S, Berg-Lyons, D, Lauber, C. L, Verduijn, W, Haasnoot, G. W, Mumm, R, Bouwmeester, H. J, Claas, F. H. J, Dicke, M, Van Loon, J. J. A, Takken, W, Knight, R, & Smallegange, R. C. (2011) Composition of human skin microbiota affects attractiveness to malaria mosquitoes. PLoS ONE 6, e28991.

[58] Chamchod, F & Britton, N. F. (2011) Analysis of a vector-bias model on malaria transmission. Bulletin of Mathematical Biology 73, 639–657.

[59] Hosack, G. R, Rossignol, P. A, & Van Den Driessche, P. (2008) The control of vector-borne disease epidemics. Journal of Theoretical Biology 255, 16–25.

[60] Kingsolver, J. G. (1987) Mosquito host choice and the epidemiology of malaria. The American Naturalist 130, 811–827.

[61] McElhany, P, Real, L. A, & Power, A. G. (1995) Vector preference and disease dynamics: A study of barley yellow dwarf virus. Ecology 76, 444–457.

[62] Roosien, B. K, Gomulkiewicz, R, Ingwell, L. L, Bosque-Perez, N. A, Rajabaskar, D, & Eigenbrode, S. D. (2013) Conditional vector preference aids the spread of plant pathogens: Results from a model. Environmental Entomology 42, 1299–1308.

[63] Sisterson, M. S. (2008) Effects of insect-vector preference for healthy or infected plants on pathogen spread: Insights from a model. Journal of Economic Entomology 101, 1–8.

[64] Thapa, I & Ghersi, D. (2023) Modeling preferential attraction to infected hosts in vector-borne diseases. Frontiers in Public Health 11, 1276029.

[65] Busula, A. O, Takken, W, De Boer, J. G, Mukabana, W. R, & Verhulst, N. O. (2017) Variation in host preferences of malaria mosquitoes is mediated by skin bacterial volatiles. Medical and Veterinary Entomology 31, 320–326.

[66] Lucas-Barbosa, D, DeGennaro, M, Mathis, A, & Verhulst, N. O. (2022) Skin bacterial volatiles: Propelling the future of vector control. Trends in Parasitology 38, 15–22.

[67] Ramırez, M, Ortiz, M. I, Guerenstein, P, & Molina, J. (2020) Novel repellents for the blood-sucking insects *Rhodnius prolixus* and *Triatoma infestans*, vectors of Chagas disease. Parasites & Vectors 13, 142.

[68] Chappell, C. R, Dhami, M. K, Bitter, M. C, Czech, L, Herrera Paredes, S, Barrie, F. B, Calderon, Y, Eritano, K, Golden, L.-A, Hekmat-Scafe, D, Hsu, V, Kieschnick, C, Malladi, S, Rush, N, & Fukami, T. (2022) Wide-ranging consequences of priority effects governed by an overarching factor. eLife 11, e79647.

[69] Tucker, C. M & Fukami, T. (2014) Environmental variability counteracts priority effects to facilitate species coexistence: Evidence from nectar microbes. Proceedings of the Royal Society B: Biological Sciences 281, 20132637.

[70] Vannette, R. L & Fukami, T. (2017) Dispersal enhances beta diversity in nectar microbes. Ecology Letters 20, 901–910.

[71] Vannette, R. L, McMunn, M. S, Hall, G. W, Mueller, T. G, Munkres, I, & Perry, D. (2021) Culturable bacteria are more common than fungi in floral nectar and are more easily dispersed by thrips, a ubiquitous flower visitor. FEMS Microbiology Ecology 97, fiab150.

[72] Belisle, M, Peay, K. G, & Fukami, T. (2012) Flowers as islands: Spatial distribution of nectar-inhabiting microfungi among plants of *Mimulus aurantiacus*, a hummingbird-pollinated shrub. Microbial Ecology 63, 711–718.

[73] Good, A. P, Gauthier, M.-P. L, Vannette, R. L, & Fukami, T. (2014) Honey bees avoid nectar colonized by three bacterial species, but not by a yeast species, isolated from the bee gut. PLoS ONE 9, e86494.

[74] Vannette, R. L, Gauthier, M. P. L, & Fukami, T. (2013) Nectar bacteria, but not yeast, weaken a plant–pollinator mutualism. Proceedings of the Royal Society B: Biological Sciences 280, 20122601.

[75] Toju, H, Vannette, R. L, Gauthier, M. P. L, Dhami, M. K, & Fukami, T. (2018) Priority effects can persist across floral generations in nectar microbial metacommunities. Oikos 127, 345–352.

[76] Vannette, R. L & Fukami, T. (2016) Nectar microbes can reduce secondary metabolites in nectar and alter effects on nectar consumption by pollinators. Ecology 97, 1410–1419.

[77] MacArthur, R & Levins, R. (1967) The limiting similarity, convergence, and divergence of coexisting species. The American Naturalist 101, 377–385.

[78] Chesson, P. L. (2000) Mechanisms of maintenance of species diversity. Annual Review of Ecology and Systematics 31, 343–366.

[79] Grainger, T. N, Letten, A. D, Gilbert, B, & Fukami, T. (2019) Applying modern coexistence theory to priority effects. Proceedings of the National Academy of Sciences of the United States of America 116, 6205–6210.

[80] Klausmeier, C. A & Tilman, D. (2002) in Competition and Coexistence, Ecological Studies, eds. Sommer, U & Worm, B. (Springer Berlin Heidelberg, Berlin, Heidelberg) No. 161, pp. 43–78.

[81] Peay, K. G, Belisle, M, & Fukami, T. (2012) Phylogenetic relatedness predicts priority effects in nectar yeast communities. Proceedings of the Royal Society B: Biological Sciences 279, 749–758.

[82] Chesson, P. L & Warner, R. R. (1981) Environmental variability promotes coexistence in lottery competitive systems. The American Naturalist 117, 923–943.

[83] Hubbell, S. P. (2001) The Unified Neutral Theory of Biodiversity and Biogeography, Monographs in Population Biology. (Princeton University Press, Princeton).

[84] Melbourne, B. A, Cornell, H. V, Davies, K. F, Dugaw, C. J, Elmendorf, S, Freestone, A. L, Hall, R. J, Harrison, S, Hastings, A, Holland, M, Holyoak, M, Lambrinos, J, Moore, K, & Yokomizo, H. (2007) Invasion in a heterogeneous world: Resistance, coexistence or hostile takeover? Ecology Letters 10, 77–94.

[85] Gill, F. B. (1988) Trapline foraging by hermit hummingbirds: Competition for an undefended, renewable resource. Ecology 69, 1933–1942.

[86] Stiles, F. G. (1975) Ecology, flowering phenology, and hummingbird pollination of some Costa Rican *Heliconia* species. Ecology 56, 285–301.

[87] Duncan, R. S & Chapman, C. A. (1999) Seed dispersal and potential forest succession in abandoned agriculture in tropical Africa. Ecological Applications 9, 998–1008.

[88] Ingle, N. R. (2003) Seed dispersal by wind, birds, and bats between Philippine montane rainforest and successional vegetation. Oecologia 134, 251–261.

[89] Ricklefs, R. E, Dodge Gray, J, Latta, S. C, & Svensson-Coelho, M. (2011) Distribution anomalies in avian haemosporidian parasites in the southern Lesser Antilles. Journal of Avian Biology 42, 570–584.

[90] Ferrari, C, Salle, R, Callemeyn-Torre, N, Jovelin, R, Cutter, A. D, & Braendle, C. (2017) Ephemeral-habitat colonization and neotropical species richness of *Caenorhabditis* nematodes. BMC Ecology 17, 43.

[91] Gillespie, D. T. (1977) Exact stochastic simulation of coupled chemical reactions. The Journal of Physical Chemistry 81, 2340–2361.

[92] Grainger, T. N, Levine, J. M, & Gilbert, B. (2019) The invasion criterion: A common currency for ecological research. Trends in Ecology & Evolution 34, 925–935.

[93] Wolfram Research, I. (2026) Mathematica, Version 15.0.

[94] R Core Team. (2025) R: A Language and Environment for Statistical Computing.

