## Supporting Information for "Regional species coexistence despite local priority effects: the overlooked role of dispersal–community feedback"

### *Full model description*

This section gives the complete model behind the Materials and Methods. Parameters and their default values are in Figure 1D.

#### *States and transitions*

A plant has $N$ flowers, each uncolonized ($\varnothing$), yeast-dominated ($Y$), or bacteria-dominated ($B$), with $\varnothing = N - Y - B$, and $P$ pollinators. Competition within a flower is fast relative to the other processes, so a colonized flower stays dominated by its first colonist until it senesces. Flowers senesce at rate $d$ and are replaced by uncolonized flowers so that the total number of flowers stays constant.

Each pollinator visits flowers at rate $c$, choosing among the plant's $N$ flowers at random. After visiting a flower in state $X$ ($X \in \{\varnothing, Y, B\}$), it leaves the plant with probability $m_X$, where $m_\varnothing = m_Y = m$; otherwise its next visit is to a random flower on the same plant. If that flower is uncolonized and the previous visit was to a yeast- or bacteria-dominated flower, the pollinator establishes yeast or bacteria with probability $e_Y$ or $e_B$. A pollinator arriving from another plant makes its first visit to a random flower on the new plant, and establishes a microbe in the same way, according to the last flower it visited on the plant it left. Bacteria also reach flowers without pollinators, carried by thrips: each bacteria-dominated flower sends bacteria to a random flower on the plant at rate $c_{B\varnothing}$, which colonize it if it is uncolonized.

In the local-scale within-plant dynamics model, we treat $\langle P\varnothing \rangle_R$, $\langle PY \rangle_R$, and $\langle PB \rangle_R$, the means of $P\varnothing$, $PY$, and $PB$ across plants, as parameters describing a regional pool, with $\langle P\varnothing \rangle_R + \langle PY \rangle_R + \langle PB \rangle_R = NP_R$, so that each plant is a continuous-time Markov chain in $(Y, B, P)$ whose transitions (Table S1) are the processes in Figure 1E, written as separate events of the chain. In Figure 1E, pollinator immigration (process 3) counts every arriving pollinator, and regional $Y$ and $B$ colonization (processes 5 and 6) are the arrivals that also colonize an uncolonized flower, so they are part of process 3. In Table S1, each arrival is one event: an arrival that colonizes a flower changes both $P$ and $Y$ or $B$, and $P$ immigration includes only arrivals that do not colonize, so its rate is that of process 3 minus those of processes 5 and 6. Figure 1E also omits senescence. For numerical solutions we blocked all pollinator arrivals at $P = P_{\max}$.

**Table S1:** Transitions of one plant with $N$ flowers in a regional pool, where $\varnothing = N - Y - B$ and $\langle PX \rangle_R$ is the mean of $PX$ across plants.

| Event | Change | Rate |
|---|---|---|
| Local $Y$ colonization | $Y \to Y + 1$ | $cP(1 - m)e_Y Y\varnothing/N^2$ |
| Local $B$ colonization | $B \to B + 1$ | $cP(1 - m_B)e_B B\varnothing/N^2 + c_{B\varnothing}B\varnothing/N$ |
| $P$ immigration | $P \to P + 1$ | $c\,[m\langle P\varnothing\rangle_R + m(1 - e_Y\varnothing/N)\langle PY\rangle_R$ $+ m_B(1 - e_B\varnothing/N)\langle PB\rangle_R]\,/N$ |
| $P$ emigration | $P \to P - 1$ | $cP(m\varnothing + mY + m_B B)/N$ |
| Regional $Y$ colonization | $P \to P + 1$, $Y \to Y + 1$ | $c\,m\,e_Y\langle PY\rangle_R\varnothing/N^2$ |
| Regional $B$ colonization | $P \to P + 1$, $B \to B + 1$ | $c\,m_B e_B\langle PB\rangle_R\varnothing/N^2$ |
| $Y$ senescence | $Y \to Y - 1$ | $dY$ |
| $B$ senescence | $B \to B - 1$ | $dB$ |

### *Reduced model*

Pollinators visit flowers much faster than flowers senesce ($c = 500$ versus $d = 0.1$ per day), so an opening left by a senescing flower is filled almost immediately so that plants have no uncolonized flowers ($B = N - Y$, $\langle P\varnothing\rangle_R = 0$). For a single opening ($\varnothing = 1$), the colonization rates in Table S1 sum to $a_Y/N$ for yeast and $a_B/N$ for bacteria, with $a_Y$ and $a_B$ as in equation 2. If $P$ does not change while the opening is filled, yeast fills it with probability $a_Y/(a_Y + a_B)$. A senescing bacteria-dominated flower is then replaced by a yeast-dominated one at rate $dB\,a_Y/(a_Y + a_B)$, and vice versa at rate $dY\,a_B/(a_Y + a_B)$, which gives the deterministic reduced model (equation 3). Pollinators arrive at rate $c(m\langle PY\rangle_R + m_B\langle PB\rangle_R)/N$ and leave at rate $cP(mY + m_B B)/N$, so for a given $Y$ they equilibrate at $\hat{P}(Y)$ (equation 1). In Figure 2B–D, arriving pollinators do not carry microbes: $m\langle PY\rangle_R$ and $m_B\langle PB\rangle_R$ are dropped from $a_Y$ and $a_B$, but not from pollinator arrivals.

With pollinator numbers fixed and no movement between plants ($m = m_B = 0$), $a_Y = c\,e_Y PY/N$ and $a_B = c\,e_B PB/N + c_{B\varnothing}B$, so

$$\frac{\mathrm{d}Y}{\mathrm{d}t} = \frac{dYB}{a_Y + a_B}\left[\frac{c(e_Y - e_B)}{N}P - c_{B\varnothing}\right]. \tag{S1}$$

$Y$ increases to $N$ if $P > P_{\text{crit}} = c_{B\varnothing}N/[c(e_Y - e_B)]$ and decreases to 0 if $P < P_{\text{crit}}$ (Figure 2A).

### *Stochastic reduced model and vacancy closure*

In the stochastic reduced model, the state of a plant is $Y$, a birth–death process on $0, \dots, N$ with rates

$$\lambda_y = d\,(N - y)\,Q_Y(y), \qquad \mu_y = d\,y\,Q_B(y), \tag{S2}$$

where $Q_Y(y)$ is the probability that the opening left by a senescing bacteria-dominated flower is filled by yeast, and $Q_B(y)$ the probability that the opening left by a senescing yeast-dominated flower is filled by bacteria.

Pollinators can arrive and leave while a flower is open, so we computed these probabilities from a Markov chain for the open flower. After a flower senesces, plant composition $(y, b)$ with $y + b = N - 1$ is held fixed while the opening is filled. With $p$ pollinators on the plant, the opening is filled by yeast at rate $A_p$ and by bacteria at rate $C_p$, and pollinators arrive at rate $U_p$ and leave at rate $D_p$:

$$\begin{aligned} A_p &= c\,e_Y \left[(1 - m)\,p\,y + m\langle PY\rangle_R\,\delta_p\right]/N^2 \\ C_p &= c\,e_B \left[(1 - m_B)\,p\,b + m_B\langle PB\rangle_R\,\delta_p\right]/N^2 + c_{B\varnothing}\,b/N \\ U_p &= c\,[m(1 - e_Y/N)\langle PY\rangle_R + m_B(1 - e_B/N)\langle PB\rangle_R]\,\delta_p/N \\ D_p &= c\,p\,(m + my + m_B b)/N, \end{aligned} \tag{S3}$$

which are the rates in Table S1 with $\varnothing = 1$ and $\langle P\varnothing\rangle_R = 0$; $\delta_p$ is 1 if $p < P_{\max}$ and 0 otherwise, so arrivals stop at $P_{\max}$. Let $q_p$ be the probability that yeast fills the opening, starting from $p$ pollinators. Conditioning on the first event gives

$$(A_p + C_p + U_p + D_p)\,q_p - U_p\,q_{p+1} - D_p\,q_{p-1} = A_p, \quad p = 0, \dots, P_{\max}, \tag{S4}$$

a tridiagonal linear system that we solved directly. The probability $r_p$ that bacteria fill the opening solves the same system with $C_p$ on the right-hand side. We averaged over the pollinator count at the moment of senescence, taken as $P \mid Y \sim \text{Poisson}(\hat{P}(Y))$ for the composition before the senescence, truncated at $P_{\max}$ and renormalized, with weights $w_p(y)$:

$$Q_Y(y) = \sum_{p=0}^{P_{\max}} w_p(y)\,q_p(y, N - y - 1), \qquad Q_B(y) = \sum_{p=0}^{P_{\max}} w_p(y)\,r_p(y - 1, N - y). \tag{S5}$$

We call this the vacancy closure.

The stationary distribution of $Y$ follows from detailed balance,

$$\pi(y) = \pi(0) \prod_{k=1}^{y} \frac{\lambda_{k-1}}{\mu_k}, \tag{S6}$$

normalized to sum to 1. We reconstructed the joint distribution of $Y$ and $P$ as $\pi(y, p) \propto \pi(y)\,\text{Poisson}(p; \hat{P}(y))$ for $p = 0, \dots, P_{\max}$, normalized over this range. For time series (Figure 3A,B), we simulated $Y$ with rates $\lambda_y$ and $\mu_y$, and $P$ with the pollinator arrival and departure rates of the reduced model, using the Gillespie algorithm, without truncating $P$.

### *Closed metacommunity*

In a closed metacommunity of many plants, the regional pools equal the pollinator output of the plants, and the stationary distribution of one plant is also the distribution of plant states across the metacommunity (20). For fixed $P_R$, the equilibrium pool solves the self-consistency equations

$$\langle PY\rangle_R = \langle PY\rangle = \sum_{y=0}^{N} \pi(y)\, y\, \hat{P}(y), \qquad \langle PB\rangle_R = NP_R - \langle PY\rangle_R, \tag{S7}$$

where $\pi$ and $\hat{P}$ depend on $\langle PY\rangle_R$ and $\langle PB\rangle_R$. Equation S7 has roots at $\langle PY\rangle_R = 0$ (yeast absent) and $\langle PY\rangle_R = NP_R$ (bacteria absent), and may have roots in between, where both species are present. We found roots by Newton's method.

The stationary probability distribution of flowers on a plant can be bimodal, with a yeast-dominated and a bacteria-dominated mode. We split $\pi(y, p)$ between modes by assigning each state to the local maximum reached by repeatedly moving to the neighbouring state (including diagonal neighbours) with the largest increase in probability. The total probability assigned to each mode gives the proportions of yeast- and bacteria-dominated plants, $p(Y)$ and $p(B)$, and the means within each mode give the means on each type of plant (Figure 5D).

### *Invasion criteria*

Yeast can invade a metacommunity of bacteria-dominated plants if a small pool of yeast-carrying pollinators, $\langle PY\rangle_R = \varepsilon$, produces a larger one: $R_Y = \langle PY\rangle/\varepsilon > 1$, with $\langle PB\rangle_R = NP_R - \varepsilon$. Likewise, bacteria can invade a metacommunity of yeast-dominated plants if $R_B = \langle PB\rangle/\varepsilon > 1$, with $\langle PY\rangle_R = NP_R - \varepsilon$. These are the metacommunity invasion criteria of ref. (20), which uses $\log R$.

For bacteria, we computed the invasion criterion as $R'_B = (NP_R - \langle PY\rangle)/\varepsilon$ with $\langle PY\rangle_R = NP_R - \varepsilon$. Because pollinator emigration balances immigration, $m\,\langle PY\rangle + m_B\,\langle PB\rangle = m\langle PY\rangle_R + m_B\langle PB\rangle_R$, which gives $R_B - 1 = (m/m_B)(R'_B - 1)$. $R_B$ and $R'_B$ therefore equal 1 at the same $P_R$, and Figure 6 is the same under either.

### *Full model*

To check the vacancy closure, we also solved the closed metacommunity with the full model (Table S1), which tracks $Y$, $B$, and $P$ without assuming that openings are refilled immediately. Its states are all $(Y, B, P)$ with $Y + B \le N$ and $P \le P_{\max}$ (17,238 states for $N = 50$ and $P_{\max} = 12$). We computed the stationary distribution by solving the linear system for the transition-rate matrix directly. The closed metacommunity then requires $\langle PY\rangle_R = \langle PY\rangle$ and $\langle PB\rangle_R = \langle PB\rangle$, with $\langle P\varnothing\rangle_R = NP_R - \langle PY\rangle_R - \langle PB\rangle_R$. For the invasion criteria, we first found the resident's equilibrium pool from the model with the invader absent (for yeast invading, the chain in $(B, P)$ with $Y = 0$ and $\langle PY \rangle_R = 0$), then computed the invader's output with its pool set to $\varepsilon$. The results of the full model were almost identical to the reduced model (see below).

### *Numerical accuracy*

We checked the invasion thresholds against three numerical choices: the invader's starting pool, $\varepsilon$; the pollinator cutoff, $P_{\max}$; and the vacancy closure, by comparison with the full model (Table S2). The values of $P_R$ at which $R_Y = 1$ and $R_B = 1$ changed by less than $10^{-5}$ across $\varepsilon = 10^{-5}$ to $10^{-9}$, and by less than $2 \times 10^{-5}$ when $P_{\max}$ increased from 12 to 18. The vacancy closure shifted the thresholds by 0.001 for yeast invasion and 0.03 for bacteria invasion, relative to the full model. The reduced model also reproduced the closed-metacommunity distributions of the full model for $0.8 \leq P_R \leq 2.8$. The full model had 0.3–0.6 uncolonized flowers per plant; relative to the reduced model, it changed the proportion of yeast-dominated plants by at most 0.014, mean pollinator numbers on either plant type by at most 0.04, and the mean number of yeast-dominated flowers on either plant type by at most 0.3. The magnitudes of $R_Y$ and $R_B$ were not converged in $\varepsilon$ (e.g., at $P_R = 1.8$, $R_Y = 2.7 \times 10^7$ at $\varepsilon = 10^{-5}$ and $7.4 \times 10^7$ at $\varepsilon = 10^{-9}$), because the invader is not rare in the stationary distribution at these $\varepsilon$; we therefore report only whether each exceeds 1. At the coexistence equilibria of the reduced model ($0.8 \leq P_R \leq 2.8$), the stationary probability of $P = P_{\max} = 12$ was at most $4.2 \times 10^{-5}$, and the equilibrium pools changed by at most 0.002% when $P_{\max}$ increased to 18 (0.02% in the full model).

**Table S2:** Values of $P_R$ at which each species' invasion criterion equals 1. Parameters as in Methods.

| Model | Numerical setting | $R_Y = 1$ | $R_B = 1$ |
|---|---|---|---|
| Reduced (vacancy closure) | $\varepsilon = 10^{-5}$, $P_{\max} = 12$ | 0.730855 | 2.895810 |
| | $\varepsilon = 10^{-7}$, $P_{\max} = 12$ | 0.730855 | 2.895811 |
| | $\varepsilon = 10^{-9}$, $P_{\max} = 12$ | 0.730855 | 2.895811 |
| Full | $\varepsilon = 10^{-5}$ to $10^{-7}$, $P_{\max} = 12$ | 0.729677 | 2.867489 |
| | $\varepsilon = 10^{-5}$ to $10^{-7}$, $P_{\max} = 18$ | 0.729670 | 2.867473 |

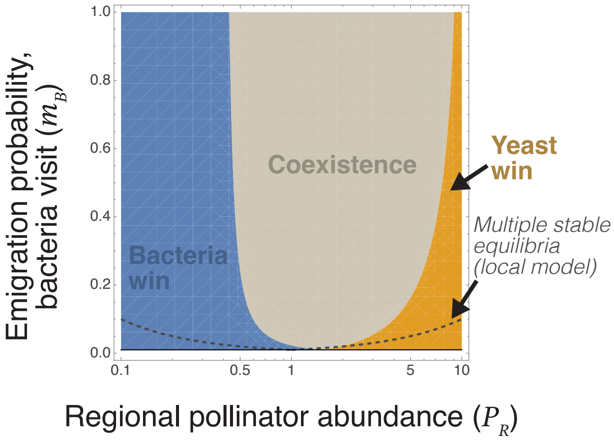


**Figure S1:** Outcome of competition in relation to regional pollinator abundance, $P_R$, and the emigration probability from bacteria-dominated flowers, $m_B$, which sets the strength of the microbe–pollinator feedback, for $m_B \leq 1$. Fill indicates outcomes, which are from invasion criteria (Methods). The horizontal line at the bottom of each pane is $m_B = m = 0.01$ (no pollinator preference). Dashed curves bound the region where a plant has alternative stable equilibria in the deterministic model without microbe immigration, $m_B/m > \max(P_R/P_{\text{crit}}, P_{\text{crit}}/P_R)$. Alternative stable equilibria are neither sufficient nor strictly necessary for regional coexistence, but the coexistence region lies largely within the region of multiple stable equilibria.
